# Idealized model of the developing visual cortex

**DOI:** 10.1101/2020.04.29.067942

**Authors:** Jennifer Crodelle, David W. McLaughlin

## Abstract

Recent experiments in the developing mammalian visual cortex have revealed that gap junctions couple excitatory cells and potentially influence the formation of chemical synapses. Though gap junctions between inhibitory cells are ubiquitous in the adult cortex, and their presence has been shown to promote synchronous network firing, their function among excitatory, pyramidal cells remains poorly understood. During development, pyramidal cells that were derived from the same progenitor cell, called sister cells, are preferentially connected by a gap junction during the first postnatal week, while chemical synapses are still being formed. Additionally, these sister cells tend to share an orientation preference and a chemical synapse in the adult cortex, a property that is diminished when gap junctions are blocked. In this work, we construct an idealized model of the mouse visual cortex during the first two postnatal weeks of development to analyze the response properties of gap-junction-coupled cells and their effect on synaptic plasticity. Further, as an application of this model, we investigate the interplay of gap-junction coupling and synaptic plasticity on the order, or organization, of the resulting cortical map of orientation preference.

**Author summary:** Gap junctions, or sites of direct electrical connections between neurons, have a significant presence in the cortex, both during development and in adulthood. Their primary function during either of these periods, however, is still poorly understood. In the adult cortex, gap junctions between local, inhibitory neurons have been shown to promote synchronous firing, a network characteristic thought to be important for learning, attention, and memory. During development, gap junctions between excitatory, pyramidal cells, have been conjectured to play a role in synaptic plasticity and the formation of cortical circuits. In the visual cortex, where neurons exhibit tuned responses to properties of visual input such as orientation and direction, recent experiments show that excitatory cells are coupled by gap junctions during the first postnatal week and are replaced by chemical synapses during the second week. In this work, we explore the possible contribution of gap-junction coupling during development to the formation of chemical synapses both into the visual cortex from the thalamus and within the visual cortex between cortical cells. Specifically, within a mathematical model of the visual cortex during development, we identify the response properties of gap-junction-coupled cells and their influence on the formation of the cortical map of orientation preference.

## Introduction

Gap junctions (GJs), or sites of direct electrical coupling between neurons, are present in the primary visual cortex (V1) at many stages of life, from infant to adulthood. In the adult cortex, gap-junction coupling among local, inhibitory cells has been shown to promote synchrony, which has been hypothesized to be important in many cognitive processes such as learning and memory [1, 2]. Though GJs have been measured between excitatory, pyramidal neurons in the adult cortex [3], there are very few experiments and the couplings were found to be very rare; consequently, their function remains unclear. Recent experiments show that pyramidal cells are coupled by GJs during the first postnatal week of development, a time at which chemical synapses are highly plastic and are just beginning to develop, leading to a question about a potential relationship between GJ coupling and the development of V1 neuron response properties.

One example of a neuron response property in the primary visual cortex is orientation preference (OP), where neurons preferentially respond to the orientation angle of a visual stimulus. In some higher-level mammals such as monkeys and cats, the visual cortex contains an ordered map of the orientation preference of each neuron, where cells preferring similar angles reside close to one another. In rodents, however, the map of orientation preference appears random and disordered, with little correlation between preferred orientation and location in cortical space.

This disordered OP map forms early in development and is dependent upon the synaptic connections from a region of the thalamus called the Lateral Geniculate Nucleus (LGN), which begins forming synaptic connections with the cortical V1 cells shortly before birth [4]. During the first postnatal week, pyramidal cells in V1 are lacking recurrent, or cortical-cortical, synapses; however, sister cells, or pyramidal cells that were derived from the same progenitor cell, are coupled through GJs [5, 6]. The strength and coupling probability of these GJs decreases steadily during the first postnatal week such that no couplings are detected in the second postnatal week [7].

During the second postnatal week, GJs between sister cells in V1 disappear and synapses begin to form between all cortical cells. Specifically, glutamatergic synapses form among the pyramidal cells, while GABAergic synapses begin to form among inhibitory, fast-spiking (FS) interneurons. In addition to the GABAergic synapses, FS cells also develop GJ coupling beginning in the second postnatal week and increasing in strength over time [8]. Turning to the map of orientation preference, visual input is not necessary for cortical cells to develop OP [9, 16]. Instead, spontaneous activity in the cortex is generated from intra-cortical circuits, as well as input from spontaneous retinal waves [10], and drives synaptic plasticity during the first two postnatal weeks [11]. By the end of the second postnatal week, a weak OP map has already developed and becomes further stabilized by visual input through the newly-opened eyes.

Our aim in this work is to better understand how the developmental timeline, including GJ-coupling among sister cells, might affect the formation of a random or disordered OP map. We develop an idealized model as a conceptual realization of a local patch of V1 during the first two postnatal weeks of development. Our model follows the set-up of Ref. [12] and the model timeline of Ref. [11], but with significant adjustments and parameters appropriate for V1. In particular, our model includes spike timing-dependent plasticity (STDP) of the feedforward synapses from LGN to V1 during the first postnatal week, as well as plasticity of the cortical-cortical recurrent excitatory synapses within V1 during the second postnatal week. Using this model, we reproduce experimentally-measured properties of GJ-coupled sister cells, such as a shared OP and preferential synaptic connectivity, and demonstrate that, during the first postnatal week, the OP of GJ-coupled cells develops faster than the OP of those cells that were not GJ-coupled. This increased learning rate results in more selectivity of the sister cells than non-sister cells at a time when synapses within V1 are beginning to form, proposing a mechanism for the “salt-and-pepper” random OP map observed in mice. We also identify mechanisms by which this OP map can become ordered as observed in higher-level mammals, further supporting our proposed mechanism for the development of disordered OP maps.

## Methods and Models

### Broad overview of experiments that measure connections between sister cells

Gap junctions between excitatory cells in the developing visual cortex of mice have only recently been discovered and their properties measured. The motivation behind this discovery was in uncovering a functional column, such as the orientation hypercolumns in monkey and cat visual cortex, which seem to be lacking in mice and rats. Because of the radial unit hypothesis, a theory positing that the cortex develops as an array of cortical columns due to clonally-related neurons traveling along the same glial fiber (axon of the progenitor cell), experimentalists began investigating the possible functional similarities between sister cells, or cells that stem from the same progenitor cell. Despite the seemingly random lateral (within layer) distribution of OPs in the visual cortex of mice (salt-and-pepper), it has been shown that radially-distributed clonally-related cells show similar stimulus feature selectivity [6], as well as preferential synaptic connectivity with fellow sister cells [5]. Each of these characteristics, measured in the adult cortex, relies on gap-junction coupling between sister cells during the first postnatal week [7, 13].

In particular, experiments by Yu et al. show that by injecting a retrovirus into progenitor cells during embryonic stages, one can illuminate a handful of its offspring (sister cells) later in development [5, 7]. To address the rate of synaptic connectivity among sister cells during postnatal development, Yu et al. concentrated on cells that were labeled by the virus to be sister cells and whose cell bodies were radially-aligned (columnar structure). They specifically measured from clusters of radially-aligned sister cells that were isolated into columns of a tangential width of about 100 *μ*m to be confident that they could distinguish between sister cells from different progenitors.Their results found that no synaptic couplings were detected between any excitatory cells, sister or non-sister, during the first postnatal week (P0-P6). In the second postnatal week, however, sister cells were measured to be coupled with an average probability of 36%, while neighboring (also radially-aligned) non-sister cells were found to be coupled with an average probability of about 6.3% (averaged over P10 to P17) [5]. The black lines in Fig 1A show the average percentage of synaptically-coupled sister cells during the second postnatal week.

**Fig 1.**
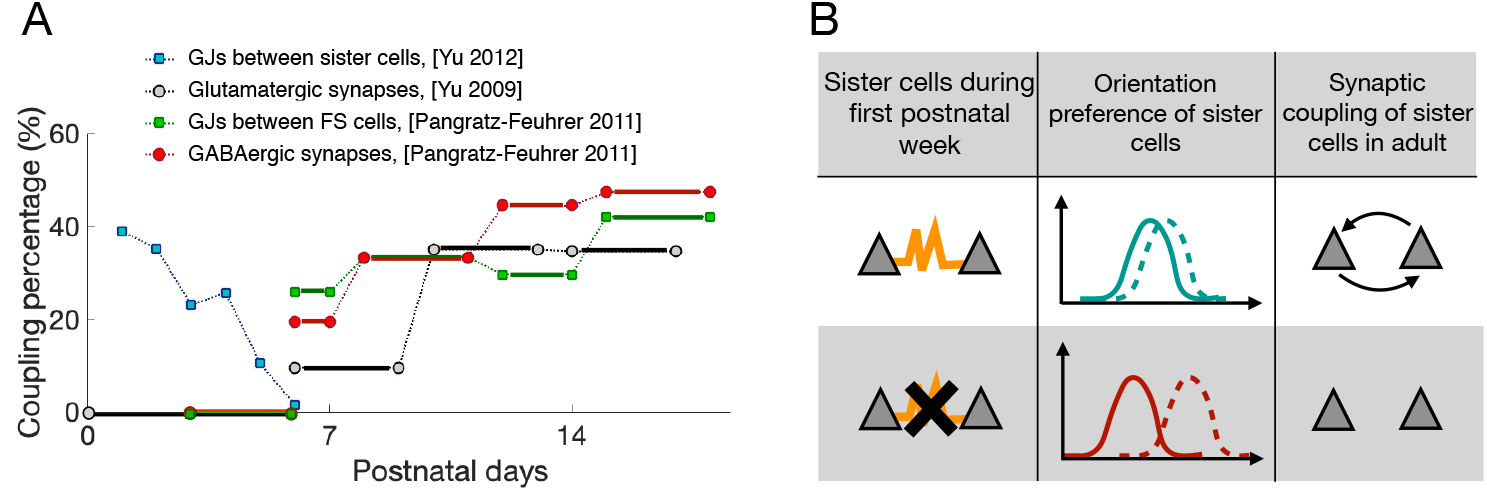
Summary of the changes in synaptic and electric coupling over the first two postnatal weeks. **A**: Plot of the different percentages of couplings over the first two postnatal weeks. The coupling probability for glutamatergic synapses is measured for radially-aligned sister cells only, while the GJs and GABAergic synapses are measured between fast-spiking (FS) cells. The experimental data was often reported as an average over several days, represented here as horizontal lines. The reference for each type of coupling is given in the legend. **B**: An illustration of the effect of GJ blocking on synapse formation among sister cells and orientation preference development. The yellow curves represent GJ coupling while the arrows represent glutamatergic synaptic coupling.

A follow-up study conducted by the same group of experimentalists used quadruple whole-cell recordings to show that these radially-aligned sister cells are coupled by GJs during the first postnatal week when chemical synapses are absent [7]. Specifically, they showed that the sister cells are preferentially coupled by GJs during the first postnatal week (28.2% for sister cells compared to 2.6% for non-sister cells, averaged over P1 to P6), with the probability of GJ connectivity decreasing steadily over the course of the first week (38.9% at P1 to ~10% at P6), see blue lines in Fig 1A. The strength of this GJ connection, as measured by the coupling coefficient (ratio of the amplitude of the response in the coupled cell to the response in the injected cell) was 5.7% for sister cells and 1.2% for non-sister cells (averaged over P1 to P6). This strength also decreases over the course of the first week for sister cells (from 7.4% at P1 to 2.3% at P6) [7].

Among inhibitory cells, GABAergic synapses and GJs form simultaneously beginning at the start of the second postnatal week [8] (in contrast to the pyramidal cells where GJs precede chemical synapses). Specifically, no GABAergic synapses or GJs are detected between FS cells from P3-P5, with the exception that one functional GABAergic synapse (out of 13 tested pairs) was detected at P5 [8] No recordings were performed before P3. Therefore, we determine that both GJ and synaptic coupling among FS cells are absent during the first postnatal week and grow during the second postnatal week, as shown by the red and green lines in Fig 1A for synaptic and GJ coupling, respectively.

To measure the effect of GJ coupling on synapse formation during development, experimentalists block the hemi-channel that connects the interiors of the two GJ-coupled cells at various timepoints. In particular, for pyramidal neurons, experimentalists show that the protein Connexin26 (Cx26) is most abundantly expressed in the neonatal cortex [14] and that blocking this protein (essentially closing the channel) largely eliminates electrical coupling among sister cells (reducing the average probability over the first week from 26% in wildtype mice to 9.8% in the Cx26-blocked mice) [7]. To test if GJ coupling might be responsible for the preferential synaptic coupling that occurs during the second postnatal week, Yu et al. repeated their earlier experiment of measuring synaptic coupling among sister cells for the case of Cx26-blocked mice as well. They found that the synaptic coupling probability dropped from the typical 30-35% between sister cells in wildtype mice to 8.2% between sister cells in Cx26-blocked mice (averaged over P10-P21) [7], demonstrating that GJ-coupling during the first postnatal week is critical to the correct circuit formation in adults.

Relatedly, experiments show that excitatory cells that share a similar orientation preference (OP) have an increased likelihood to also be synaptically coupled [15]. Since GJ coupling during the first postnatal week is necessary for preferential synaptic coupling in adults, several experimentalists set out to assess the role that GJ coupling might play in stimulus feature selectivity, such as orientation preference [13, 17]. Li et al. used retrovirus labeling (similar to Yu et al.) to show that radially-aligned sister cells in L2/3 of mouse visual cortex have a similar OP. Specifically, they showed that about 59% of measured sister cells have similar OPs (difference in preferred angle less than 30^*◦*^), while neighboring non-sister cells exhibit a difference in OP distribution that was not significantly different from the uniform distribution [13]. When a GJ blocker was employed, the effect was destroyed; the distribution of OP difference for sister cells was no longer significantly different from the uniform distribution or the non-sister cell distribution. Figure 1B shows a schematic of the effect of GJ blocking between sister cells in the first postnatal week.

As for the organization of sister cells in the postnatal cortex, sister cells are derived from radial glial cells and migrate to their end location by traveling down the axon of the glial cell. While traveling, these sister cells begin dispersing laterally such that by the end of the second postnatal week, they are dispersed up to 500 *μ*m in radius (see Fig S1 in [5]). Then, sister cells become sparsely intermingled in the mouse visual cortex, with sister cells outnumbered by non-sister cells in a local volume (100-500 *μ*m in diameter) by a factor of six [18, 19], a property that seems to be essential for proper synaptic development [20]. While previous experiments concentrated on small groups of radially-aligned sister cells within a radius of about 100-120 *μ*m [6], Ohtsuki et al. used a gene-targeting system to label all of the offspring of a single cortical progenitor cell (about 600 neurons). They showed that sister cells are typically distributed throughout layers 2-6, with the clusters of sister cells having a tangential diameter of about 300-500 *μ*m [17]. Measuring the OP of all sister cells, they found that about 50% of each sister-cell pair tends to have an OP within 40^*◦*^. They reason that this probability is smaller than that measured by Li et al. due to the significantly larger population of cells that they are measuring.

In summary, these experiments show that, during the first postnatal week, radially-aligned sister cells in the cortex are coupled by GJs, but contain no recurrent synaptic connections. At the start of the second postnatal week, GJs between sister cells disappear and chemical synapses among inhibitory cells (together with GJs between inhibitory cells) and synapses among excitatory cells begin to form. Recall Fig. 1B for the percentages over the two postnatal weeks. As concerns the effect of GJ-coupling between sister cells, experiments show that sister cells that were coupled by a GJ during the first postnatal week preferentially develop a chemical synapse during the second postnatal week and are more likely to have a similar OP by the time of eye-opening.

### The Mathematical Model

In this section, we describe our idealized model of development. Specifically, we use a similar framework as the model in Ref. [12] for the study of spike-timing-dependent plasticity (STDP), but incorporate a more realistic and stable STDP learning rule for the visual cortex adapted from Refs. [22, 25] with added inhibitory plasticity as in Ref. [28]. The details are as follows.

We consider 1000 feedforward synapses, representing input from LGN to the visual cortex, coupled to our model neuronal network of either 400 or 256 cortical cells. The cortical neurons are randomly assigned to be excitatory with 80% probability or inhibitory with 20% probability. The subthreshold voltage of the *i*th cortical neuron of type *Q* = {*E, I*} is described using the leaky integrate-and-fire equation as follows

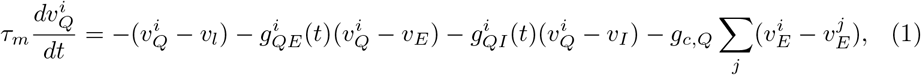

where *τ*_*m*_ = 20 ms, *v*_*l*_ = −60 mV, *v*_*E*_ = 0 mV, and *v*_*I*_ = −80 mV. Once the voltage reaches a threshold of −45 mV, the neuron is said to have spiked, the spike time is recorded, and the voltage is reset to −60 mV. Gap junctions are included only among excitatory neurons, such that the conductance term *g*_*c,Q*_ takes on a nonzero value *g*_*c*_ for *Q* = *E* and zero for *Q* = *I*, and are incorporated into the model through a direct resistive term where 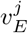 is the voltage of the *j*th pre-junctional neuron; see the last term in Eq (1). In addition, to model the spikelet induced in the post-junctional cell in response to an action potential in the pre-junctional cell, a 1 mV instantaneous jump in voltage of the post-junctional cell is included, as in previous models [11, 21].

The cortical synaptic conductances are modeled as having instantaneous rise times and exponential decay at each received spike time so that the excitatory and inhibitory conductance traces, respectively, follow the equations

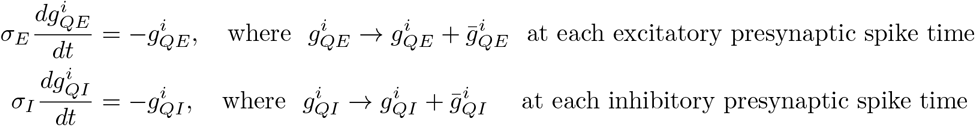

where the neuron type of the postsynaptic cell is represented by *Q* = {*E, I*}, *σ*_*E*_ = 11 ms and *σ*_*I*_ = 15 ms. Note that the synaptic conductances have been normalized by the leakage conductance and are thus unit-less. The maximal excitatory conductance strength, 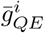, and inhibitory conductance strength, 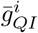, can each take one of the following values: 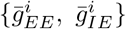 and 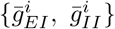 where the subscript *XY* denotes the direction of coupling from *Y* to *X*. We implement an absolute maximum on all excitatory synapses at 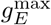 and on all inhibitory synapses at 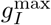. In this model, the conductances 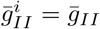 and 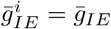 are held constant at 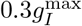 and 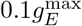, respectively, for all cells, while 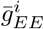 and 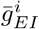 are plastic, changing with rules defined in the following subsection.

The external drive to the cortical network has two components: synaptic input from the LGN and a generic background drive to all cells. This external drive affects the excitatory conductance, 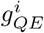, as follows

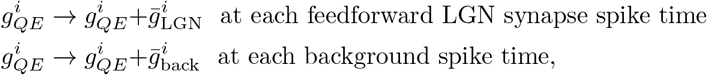

where 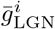 is plastic, but 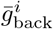 is constant at 0.02. The spike times of the background drive are generated from a Poisson process with rate 0.5 Hz. Each feedforward LGN synapse generates spikes using a Poisson spike train with a firing rate that depends on its own label. Specifically, the firing rate of LGN synapse labeled *a* in response to a stimulus at input location *s* is given by

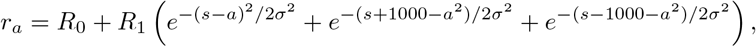

as in [12], where *R*_0_ = 5 Hz, *R*_1_ = 20 Hz, and *σ* = 80. Input to these synapses consists of brief presentations of a uniformly randomly-chosen stimulus index (*a* in above equation) for a period of time that is chosen from an exponential distribution with mean 20 ms. All cortical cells receive input from LGN synapses with a 25% probability. While the inhibitory cells have a constant LGN feedforward synaptic strength randomly chosen uniformly between 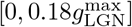, the excitatory cells contain a plastic or variable strength, 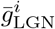.

### Plasticity Rules

In the model, feedforward LGN synapses to excitatory cortical cells, as well as the synapses between cortical excitatory cells, are plastic, with the strength of their connection, 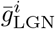 and 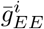, respectively, obeying the minimal triplet rule for the visual cortex [22]. We use the triplet rule rather than the standard pre-post STDP rule that was used in [12] because we wish to reproduce the realistic bi-directional coupling that develops in the visual cortex of mice, a feat that cannot be accomplished with the pair-based STDP rules due to their nature of developing only unidirectional synapses. In addition, experiments show that the STDP curves exhibited by pyramidal cells in the visual cortex of mice do not follow the typical slightly-asymmetric shape of potentiation and depression as in [23], but rather potentiation only occurs if the post-synaptic neuron had recently fired a spike of its own [22, 24].

The triplet rule is illustrated in Fig 2A and described as follows. For each pre- and post-synaptic spike, the strength of the synapse from the pre- to post-synaptic cell, 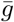, (dropping the *EE* subscipt) is updated as follows:

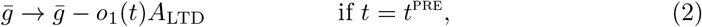

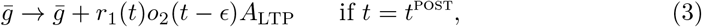

where *A*_*LT D*_ and *A*_*LT P*_ represent the strength of depression and potentiation, respectively. The tracer variables follow the equations

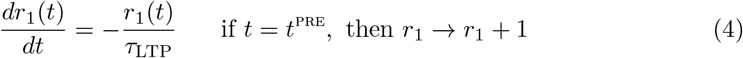

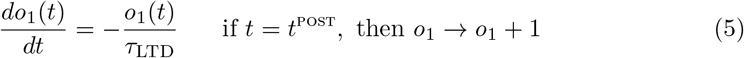

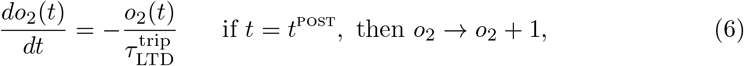

where *r*_1_(*t*) represents a pre-synaptic tracer, while *o*_1_(*t*) and *o*_2_(*t*) represent post-synaptic tracers. Note that each neuron carries its own tracer variable, but the *i* index has been dropped here for clarity. The timescales of these tracer variables are as follows: *τ*_LTP_ = 16.8 ms, *τ*_LTD_ = 33.7 ms, and 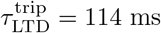. To stabilize network activity, we implement a homeostatic mechanism in the form of a rate detector that acts on a fast timescale, known to stabilize the dynamics induced by the minimal triplet rule into recurrent excitatory networks [25]. This homeostatic mechanism works by allowing the amount of depression, *A*_LTD_, to change as a function of a moving-average of the post-synaptic firing rate, 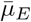:

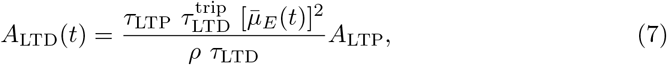

where the timescales *τ*_*LTP*_, 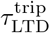, and *τ*_LTD_, are those from Eqs (4) - (6), and *ρ* is the target firing rate, chosen to be 8 Hz to replicate the low firing rate of the mouse visual cortex during early development [27]. The moving average of the firing rate, 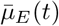, is found by taking a low-pass filter of its spike train as follows

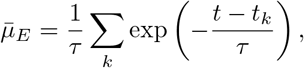

where *t*_*k*_ represents the *k*th spike time that occurred prior to the current time *t* and *τ* = 1 s. Note that the synaptic strength 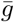 in Eqs (2) and (3) can take on either 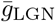 for synapses from LGN to the cortex, or 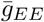 for synapses among excitatory cortical neurons. These synapses also have different learning rates, 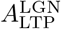 and 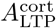, for the feedforward synapses and recurrent cortical synapses, respectively. See Table 1 for a comprehensive list of parameter values.

**Table 1.** Model parameter values parametrized for the visual cortex.

| Neuron Parameters |  |  |  |
| --- | --- | --- | --- |
| Membrane time constant, $\tau_m$ | 20 ms | Leakage reversal potential, $v_l$ | -60 mV |
| Excitatory reversal potential, $v_E$ | 0 mV | Inhibitory reversal potential, $v_I$ | -80 mV |
| GJ conductance, $g_c$ | 0.06 | Background firing rate, $\nu$ | 0.5 Hz |
| Excitatory synaptic time constant, $\sigma_E$ | 11 ms | Inhibitory synaptic time constant, $\sigma_I$ | 15 ms |
| background spike strength, $\bar{g}_{\text{back}}$ | 0.02 | | |
| Plasticity Parameters |  |  |  |
| Learning rate for LGN, $A_{\text{LTP}}^{\text{LGN}}$ | 0.005 | Time constant, $\tau_{\text{LTP}}$ | 16.8 ms |
| Learning rate for V1, $A_{\text{LTP}}^{\text{cort}}$ | 0.015 | Time constant, $\tau_{\text{LTD}}$ | 33.7 ms |
| Time constant, $\tau_{\text{LTD}}^{\text{trip}}$ | 114 ms | Maximum LGN weight, $g_{\text{LGN}}^{\text{max}}$ | 0.02 |
| Maximum I → E weight, $g_I^{\text{max}}$ | 0.05 | Max E → E weight, $g_E^{\text{max}}$ | 0.025 |
| Learning rate for iSTDP, $A_{\text{iSTDP}}$ | 0.008 | Time constant, $\tau_{\text{iSTDP}}$ | 20 ms |
| Target firing rate, $\rho$ | 8 Hz | | |

**Fig 2.**
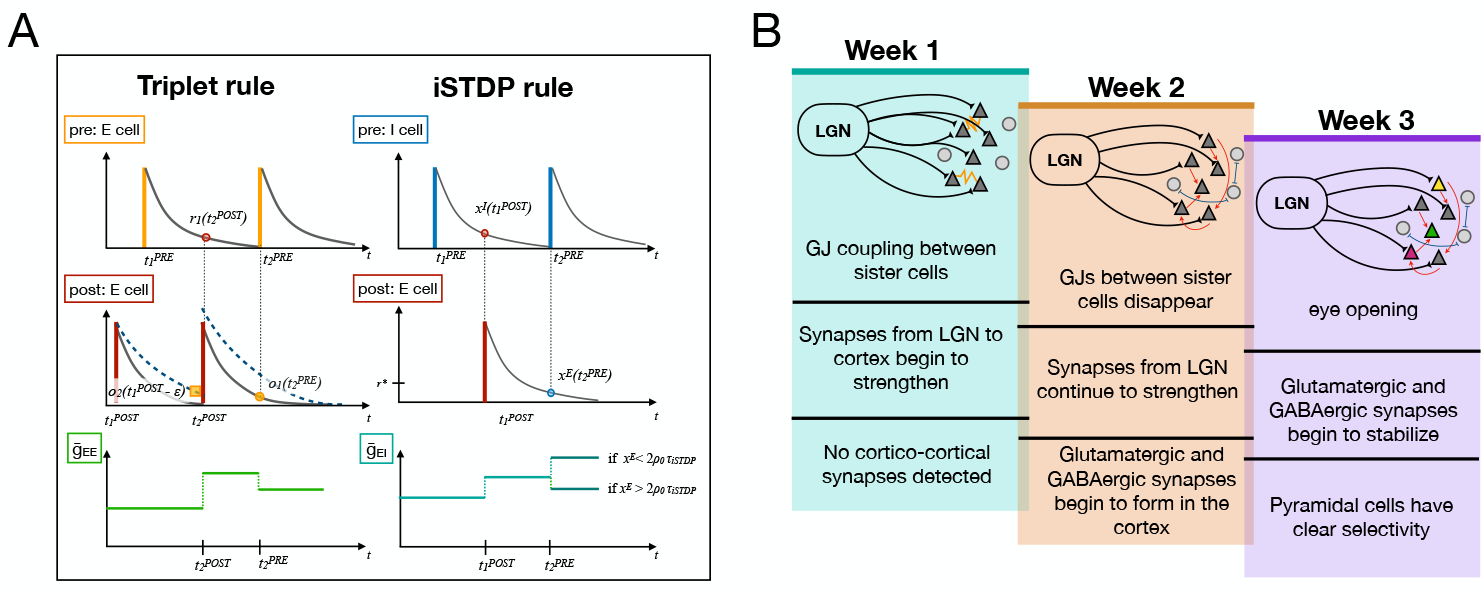
**A**: Schematic of the triplet rule for excitatory synapses and the iSTDP rule for inhibitory synapses onto excitatory neurons. For the triplet rule, an example weight update resulting from a post-pre-post interaction (at 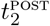) and a pre-post-pre interaction (at 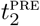 is shown. For inhibitory plasticity, the weight update is shown for one potentiation example for a spike time of the excitatory neuron (at 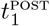) and an update that depends on the tracer variable of the postsynaptic cell, *x*^*E*^, the target firing rate *ρ*, and the timescale of the inhibitory plasticity, *τ*_iSTDP_. The vertical axis of the top two plots represent the tracer values, while the vertical axis of the bottom plots represents the synaptic conductance strength. **B**: Timeline of events and schematic of the model for the first three postnatal weeks.

In addition to the plasticity introduced on the feedforward and recurrent excitatory synapses, we include plasticity on the synapses from inhibitory neurons to excitatory neurons in the cortex [28]. The motivation behind this inhibitory plasticity is that we found it necessary for the inhibition in the network to mediate the excitation for proper development to occur. If the inhibitory synapses were constant at a high value, then the cortical cells would not fire any action potentials. However, if the inhibition was constant at a low value, then as the excitatory recurrent synapses grew, the firing rate of the network would also grow, and the network would become unstable, a common phenomena for plastic recurrent excitatory networks. Therefore, we chose to model inhibitory plasticity as a stabilizing mechanism, as has been done previously in Ref. [28].

A schematic of this inhibitory plasticity can be found in Fig 2A. The synapse from a pre-synaptic inhibitory cell to a post-synaptic excitatory cell updates according to the following rule

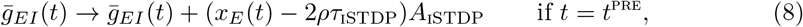

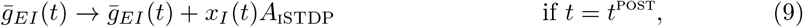

where *A*_iSTDP_ is the learning rate and *ρ* = 8 Hz is the target firing rate of the excitatory cells [the same as in Eq (7)]. Each cell has a tracer variable *x*_*Q*_ for *Q* = *E, I* that follows the equation

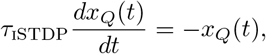

where *τ*_iSTDP_ = 20 ms and *x*_*Q*_(*t*) → *x*_*Q*_(*t*) + 1 at each spike time of the cell, similarly to the tracer variables *r*_1_*, o*_1_ and *o*_2_ in Eqs (4) - (6). Note the interpretation of these plasticity rules: when the spiking of a pre- and post-synaptic inhibitory and excitatory cell, respectively, occurs within a time window of *τ*_iSTDP_, either potentiation or depression occurs at each pre-synaptic (inhibitory) spike [as per Eq (8)], while only potentiation occurs at each post-synaptic (excitatory) spike [as per Eq (9)].

Development is simulated by connecting a subset of the cortical cells by GJs and allowing the LGN synapses onto all excitatory cortical cells to learn for a period of time (which varies in this work), simulating the first postnatal week of development (see blue panel in Fig 2B). Then, once simulation is in the second postnatal week, gap junctions are turned off [by setting *g*_*c,E*_ = 0 in Eq (1)], and recurrent synapses are turned on. Specifically, 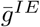 and 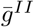 go from zero to nonzero values; 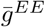 updates (and LGN synapses continue to update) according to the rules defined in Eqs (2) - (3); and 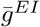 updates according to the rules defined in Eqs (8) - (9). We simulate this network until the recurrent cortical weights have stabilized and each cortical cell has developed an input preference (called the OP in this work). We note that the network operates in an asynchronous regime known to accentuate the performance of STDP [26].

Tuning properties of the cortical cells are determined by taking the final weights from the simulated network and, for each input stimulus preference from 0 to 1000 in increments of 20, we record the firing-rate responses for all neurons averaged over two seconds of simulation time. Then, tuning curves are calculated for each cortical cell by determining the firing rate of that cell for each input stimulus and normalized by the maximum firing rate across all cells. The OP of the cortical cell is determined as the stimulus location that gives the greatest response.

## Results

In this section, we describe simulation results for three realizations of the cortical network, each with progressively more realistic connectivity properties. These realizations of the network are chosen to illustrate three characteristics of synaptic development in the presence of GJs: (i) GJ coupling serves to increase the rate of synaptic learning of the projections from LGN to the GJ-coupled cells. This increased rate of learning leads pairs of GJ-connected cortical cells to develop similar OPs and preferential synaptic connections; (ii) The distribution of OP that develops when cortical synapses are allowed to form in an all-to-all fashion between excitatory cells is nearly uniform, spanning all possible OPs equally. This distribution becomes less uniform (one prominent OP emerges) when GJs are not present during the first phase of development and cortical synapses are allowed to learn earlier in the developmental timeline; (iii) The OP map that develops when cortical synapses are restricted to form within a small radius centered on each cell can range from ordered to disordered depending on the relative timing of cortical synapse development to that of LGN synapse development, as well as the presence or absence of GJ coupling between cortical cells during the first developmental phase.

### Realization (i): GJs and receptive field development

We begin by studying the development of the feedforward LGN synapses onto the cortical cells and how this learning is affected by GJ coupling. Then, we allow cortical synapses to form in an all-to-all fashion and investigate the effect of GJ-coupling on synapse formation. Specifically, we demonstrate that a GJ between cortical cells allows those cells to develop a similar OP and a preferential bidirectional synapse, as shown in experiments. In addition, we illustrate that GJ-coupled cells develop an OP at a faster rate than those that are not coupled by a GJ, leading us to hypothesize that the inclusion of GJs during the first postnatal week will enhance spatial disorder in the formation of the OP map.

To illustrate these effects, we use a 400-neuron idealized cortical network in which 20% of the cells are inhibitory and 80% are excitatory. We allow two excitatory cells to be coupled by a GJ with a 50% probability such that about half of the excitatory population is GJ-coupled in pairs (similarly to the small network explored in Ref. [11]). Following the experimental timeline (recall Fig 1A and Fig 2B), we simulate the first postnatal week of development by allowing the feedforward synapses to learn, via the rules discussed in the Methods and Model section, for 600 seconds of simulation time. During this time, about half of the excitatory cells are GJ-coupled in pairs, while the recurrent synaptic connections are set to zero (phase 1 of development). At the end of this phase, we turn off the GJs between cortical cell pairs and allow recurrent cortical synapses to learn together with the feedforward synapses from the LGN (phase 2 of development).

Due to the competitiveness of the STDP learning rule, the strength of the LGN synapses onto one cortical cell develops in such a way that about half of them potentiate to the maximum possible synaptic strength and half are depressed to zero, see Fig 3A. Further, the synapses that become potentiated tend to have a similar labeling (i.e., respond preferentially to a similar input value), resulting in an input preference for the cortical cell at the end of the simulation, see Fig 3B. This input preference is what we refer to as the OP of the cortical cell in this work. The cells that were coupled by a GJ during the first phase of development, i.e., during the time of feedforward learning, develop similar OPs, while cells that did not contain GJ coupling are not likely to share an orientation preference, see Fig 3C,D. Finally, the model reproduces the experimentally-observed behavior for GJ-coupled cells to preferentially form bidirectional synapses, see Fig 3E. Note that the probability of finding bidirectional synapses between GJ-coupled cells is much higher in the model than those observed in real cortex (26% in [7] compared to almost 80% here) since we are directly comparing GJ-coupled cells, while the experiments tested all sister cells (only a fraction of which are coupled by a GJ).

**Fig 3.**
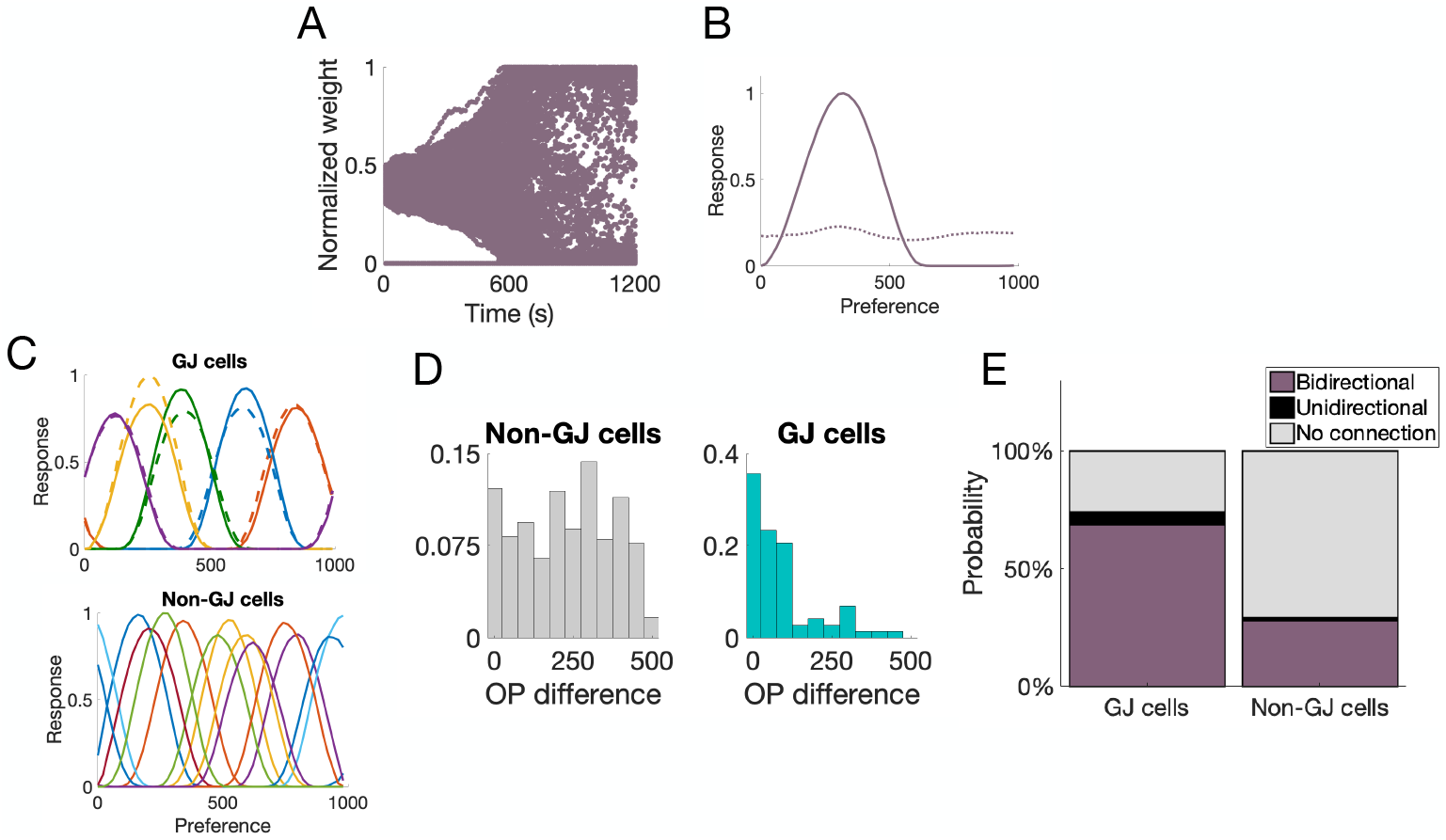
Measurements from a 400-neuron network with pairwise GJ coupling. **A**: Progression of the synaptic weights from LGN to one sample excitatory cortical cell; **B**: Tuning curve of this sample cell before (dotted curve) and after (solid curve) feedforward synaptic learning; **C**: Sample tuning curves. (top) Tuning curves of five sample GJ-coupled pairs, where matching colors indicate the GJ-coupled pairs. (bottom) tuning curves of non-GJ-coupled excitatory cells; **D**: Distribution of the difference in OP between GJ-coupled pairs (blue) and non-GJ-coupled cells (gray); **E**: Probability of a bidirectional synapse (purple), a unidirectional synapse (black), or no synapse at all (gray) between GJ-coupled cells and non-GJ-coupled cells.

We noticed in our simulations that the GJ-coupled cells tend to develop an OP much sooner than the non-GJ-coupled cells. This means that the feedforward synapses from LGN onto the GJ-coupled cells learn much faster than those synapses onto cells that are not GJ-coupled, see Fig 4A for one example cell that was GJ-coupled and one that was not. To see this effect over all GJ-coupled and non-GJ-coupled pairs, for each cell, we average together all feedforward synapses that potentiated to at least 70% of the maximum synaptic weight, and then average over all cells in each population; see Fig 4B. Notice that the slope of the average synaptic strength is much larger for those cells with GJ coupling during the first phase of development than for those cells that did not have GJ-coupling during the first phase of development.

**Fig 4.**
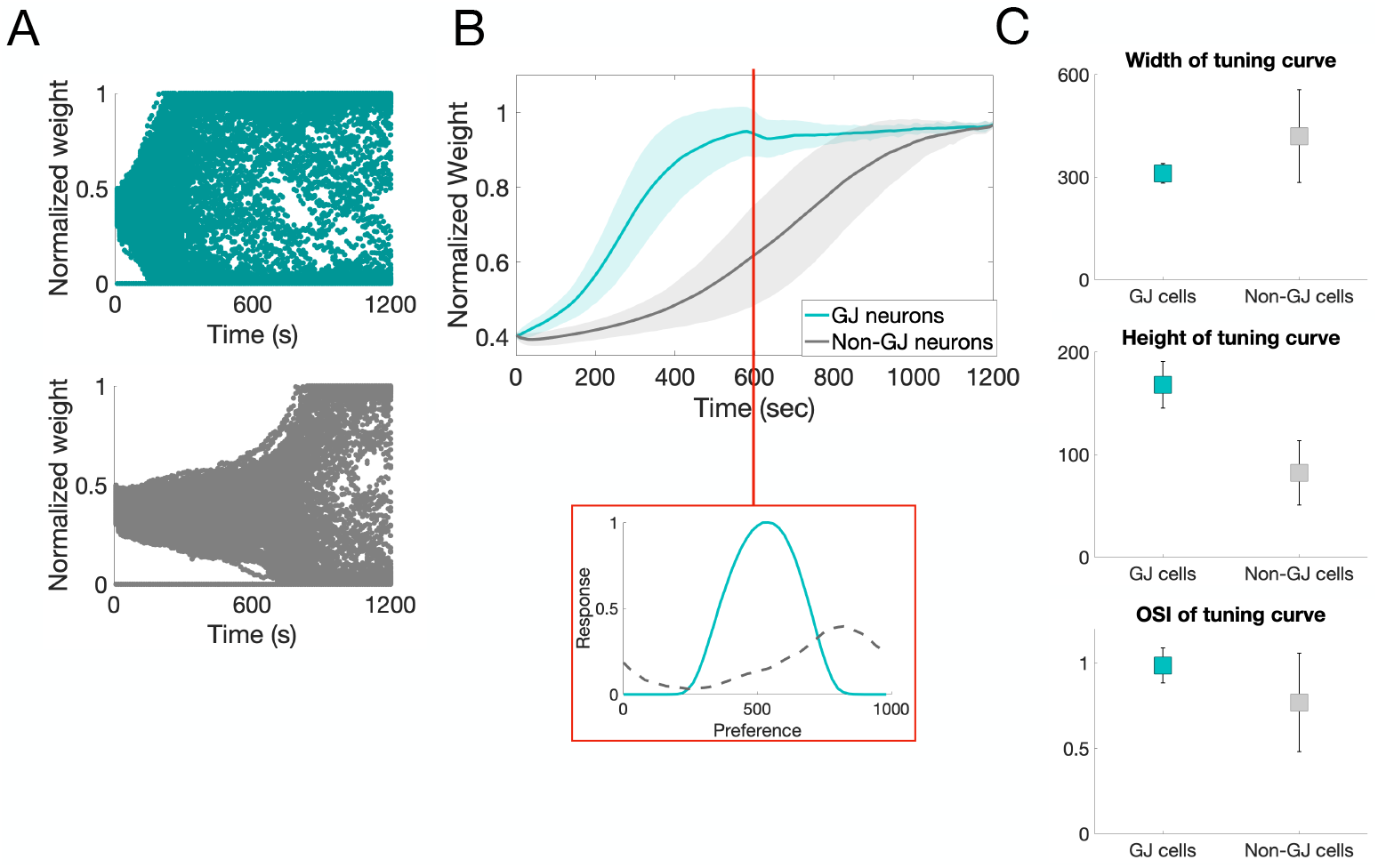
Rate of learning properties of GJ- vs. non-GJ-coupled cells. **A**: Top (bottom): Progression of the feedforward synaptic weights onto a sample GJ-coupled (non-GJ-coupled) neuron. **B**: The average weight progression (curve) and standard deviation (shaded region) for all GJ-coupled neurons (blue) and all non-GJ-coupled neurons (gray). Inset: Sample tuning curve for a GJ-coupled neuron (solid) and a non-GJ-coupled neuron (dashed) after 600 seconds of simulation time, before recurrent connections begin to form. **C**: Width and height of the tuning curves measured for all cells in the network, as well as the orientation-selectivity index (OSI), the average reported as the center of each square, the standard deviation as error bars, over all cells in each group.

The result of an increased learning rate of the LGN synapses onto the GJ-coupled cells is that the GJ-coupled cells are more selective for orientation (have more clearly-defined tuning curves) than the non-GJ-coupled cells. To investigate this hypothesis, we measure properties of the tuning curves of the GJ-coupled and non-GJ-coupled neurons at the end of the first phase of development, before cortical synapses have been allowed to learn. Figure 4B (bottom) shows the tuning curve of a GJ-coupled cell (blue) and non-GJ-coupled cell (gray) at the end of the first phase of development. Notice that the GJ-coupled cell has more selectivity than the non-GJ-coupled cell, as indicated by the tall thin peak. This effect can be quantified over all cells in the network by considering the width and height of the tuning curve for all GJ-coupled cells and non-GJ-coupled cells. In addition, we calculate the orientation-selectivity index (OSI), a measure for selectivity of a cell, defined as follows

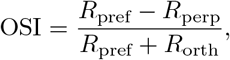

where *R*_pref_ is the firing rate of the neuron at its preferred orientation and *R*_orth_ is the firing rate of the neuron at the orthogonal orientation (in this work, the orthogonal orientation corresponds to the orientation that is 500 units away from *R*_pref_). An OSI value close to 1 indicates high selectivity and a value close to 0 indicates no selectivity. Figure 4C shows the average of these three properties over all GJ-coupled (blue) and non-GJ-coupled (gray) neurons. Notice that, for three measures of orientation selectivity, GJ-coupled neurons clearly have more selectivity than non-GJ-coupled cells at the time that cortical synapses begin to form.

The implication of GJ-coupled cells having more selectivity than non-GJ-coupled cells is that their tuning properties (i.e., their OPs) are less likely to be influenced (changed) by the recurrent cortical synapses when they begin to form during the second phase of development. If we consider sparsely-coupled GJs (e.g., pair-wise) during the first phase of development, and expect that GJ-coupled cells will preferentially develop similar OPs while different sets of GJ-coupled cells develop different OPs (since they do not communicate during the first phase), then we expect to see pairs of cells with similar OPs scattered throughout the cortex. Assuming that the GJ-coupled cells are sufficiently selective at the initiation of recurrent cortical learning such that their OP does not change during this second phase, one would expect that the final OP map in this case would be salt-and-pepper, as demonstrated in the left schematic of Fig 5. On the other hand, if GJs did not exist during the first phase of development while LGN synapses were forming, then the cortical cells would not be sufficiently selective by the time that recurrent synapses formed, and the development of the cortical recurrent synaptic connections would influence the final OP of each cell. This might result in an OP map that has order, as demonstrated in the right schematic of Fig 5. In the next section, we begin to investigate how the selectivity of the GJ-coupled cells at the start of recurrent cortical learning phase might influence the properties of the final OP map.

**Fig 5.**
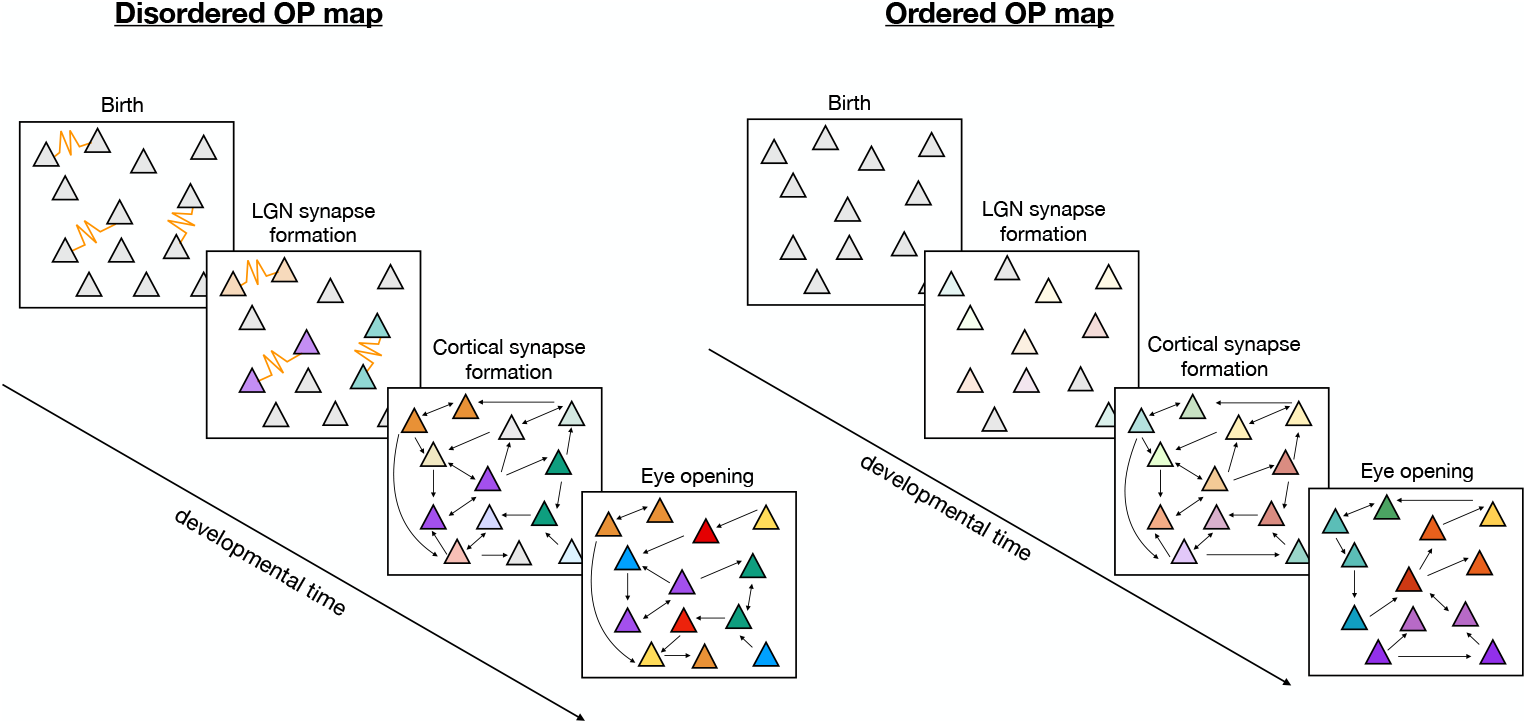
A schematic representing how GJs during the first week might lead to a disordered (i.e., salt-and-pepper) OP map. Each stage drawn here represents: (i) The first postnatal week as labeled by “Birth” and “LGN synapse formation”, (ii) the second postnatal week while chemical synapses are forming, and (iii) the resulting cortical recurrent synapses and OP as indicated by the color of the cell. Transparency represents selectivity, where opaque colors indicate a higher amount of selectivity or sharp tuning.

### Realization (ii): recurrent synapse formation (all-to-all network)

In this section, we begin to study the effect of GJ-coupling on the formation of cortical synapses by including sister-cell groups in the excitatory population and varying the time at which the synapses between cortical cells form. We do this so that we may consider more realistic GJ coupling between sister cells (rather than simply between pairs as in the previous section) in the recurrent network and investigate how the resulting OPs of the cortical cells is affected by GJ coupling during the first postnatal week. We also decrease the model network size from 400 to 256 neurons for ease of computation. To create sister-cell groups, we divide the excitatory population into six groups with equal probability, where each group represents a set of sister cells (i.e., all cells in each group are sister cells to only those cells in that group). The motivation behind choosing six groups of sister cells is that, in mouse V1, sister cells are intermingled with other sister cells and outnumbered in a local volume by a factor of six [17]. We assume that 256 neurons corresponds to a small enough volume of the cortex that we can consider only six groups of sister cells that are randomly distributed in the space. Within each sister-cell group, each neuron has a 5% probability of being coupled to a sister cell by a GJ. Figure 6A shows a count of the number of cells in each sister group along with the probability of GJ coupling in each group. Note that this coupling percentage is much sparser than the ~ 28% coupling probability measured experimentally for radially-aligned sister cells [7]. We found that is was necessary to require a sparse GJ-coupling during the first postnatal week for the GJ-coupled cells to exhibit the experimentally-measured properties of OP sharing and preferential synaptic coupling. We explore the effects of larger GJ-coupling percentages in the Supporting information.

**Fig 6.**
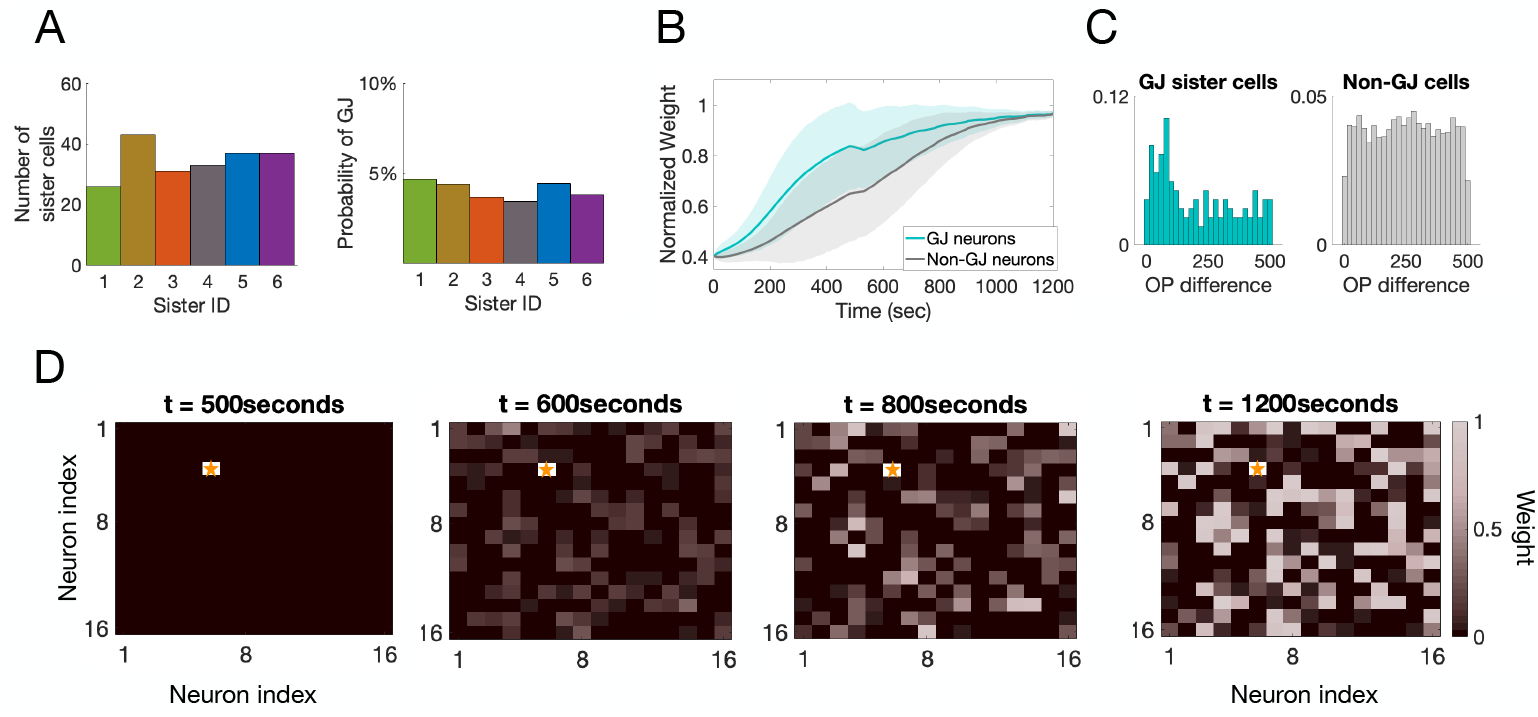
Measurements from a 256-neuron network with all-to-all potential cortical connectivity. **A**: The number of cells (left) and the probability GJ-coupling (right) within each of the six sister groups. **B**: The rate of LGN synaptic learning averaged over all GJ-coupled cells (blue) and non-GJ-coupled cells (gray). **C**: The distribution of differences in OP for GJ-coupled cells (blue) and non-GJ-coupled cells (gray). **D**: The normalized recurrent cortical weights for one example excitatory cell, indicated by a yellow star. The recurrent connections begin at *t* = 500 seconds and the entire simulation was run for 1200 seconds.

The response properties measured for the pairwise GJ-coupled 400-neuron network remain in this 256-neuron network, including the increased rate of learning for GJ-coupled cells compared to non-GJ-coupled cells, see Fig 6B, and the preference for GJ-coupled cells to share an OP, see Fig 6C. The recurrent synapses between excitatory cells can be all-to-all, but due to the competitive STDP rules, each excitatory cell forms a strong synapse with only about half of the other excitatory cells (the other half decay just as in the feedforward synapses). Figure 6D demonstrates an example of the progression of the cortical synapses onto one excitatory cortical cell, indicated by the star. Note that some of the weights correspond to inhibitory synapses and some are excitatory synapses.

We use this network to begin to test our hypothesis that GJ-coupling during the first phase of development leads to disorder in the OP map. First, we show that if the GJs are turned off during the time that LGN synapses are learning (the first phase of development), the distribution of OPs that forms has more order than the one that forms when GJs are present during the first phase of development. Recall that we do not include any spatial effects in this second realization of the model, and we allow any cell in the cortex to form a synapse with any other cell. Thus, to measure order in the OP map, we use the idea from Ref. [12] that when cortical recurrent connections are allowed to form between all cells in the network, all cortical cells develop a similar OP. Figure 7A shows that the OPs in the network without GJ coupling tend to cluster around one value (~ 375), indicating that the recurrent connections influence the resulting OP of each cell, while the network with GJ coupling during the first phase of learning has a more uniform distribution of OPs. To quantify this, we calculate the distance between each resulting OP distribution and the uniform distribution (sum of the squared difference). Notice that the network without GJ coupling during the first phase of development has a distance of 0.182, while the OP distribution for the network containing GJ coupling has a distance about half of that value at 0.099. This indicates that the inclusion of GJ coupling during the first phase of development results in an OP distribution closer to the uniform distribution, where each OP has equal likelihood of occurring.

**Fig 7.**
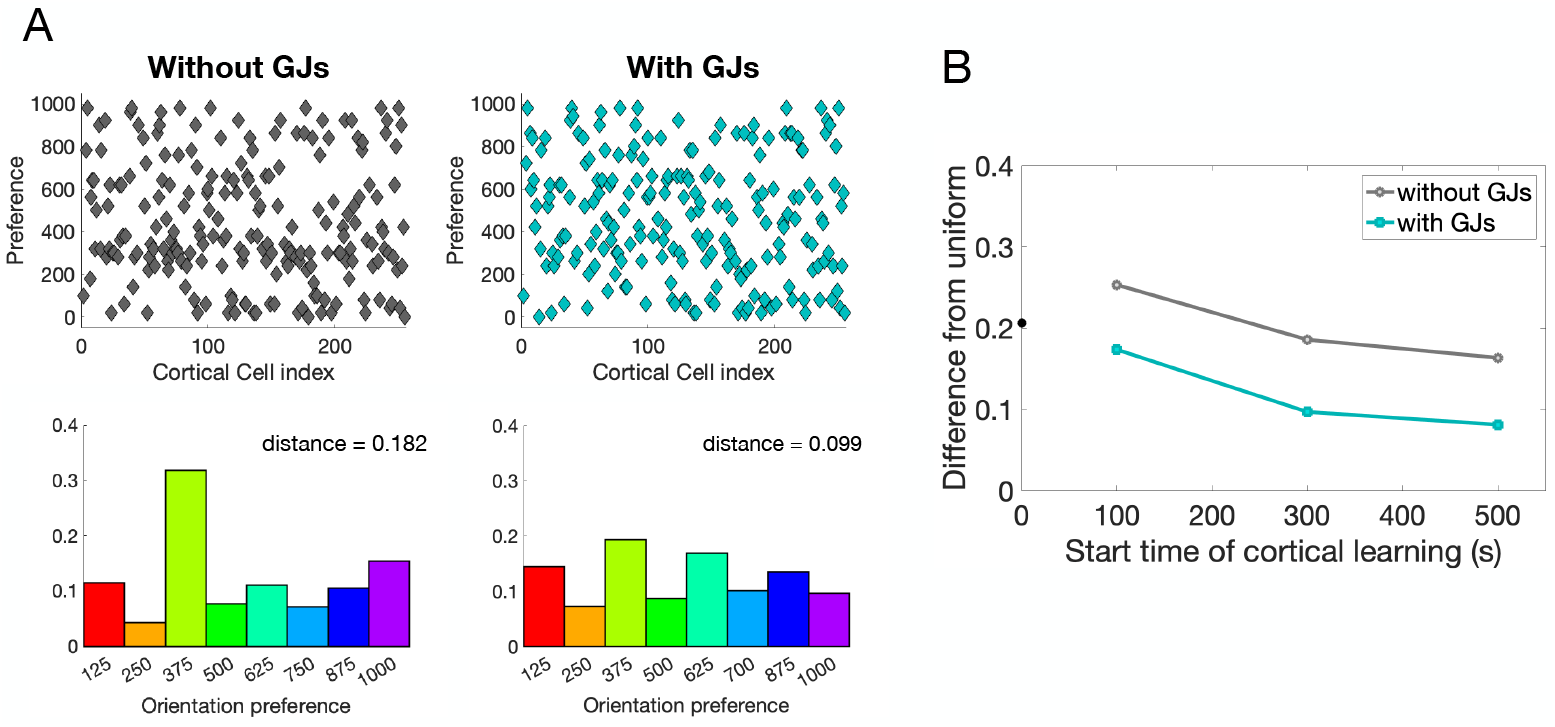
Distribution of OPs comparing a network containing GJ coupling to one without. **A**: Left (right): Top: Plot of each cell’s OP; Bottom: distribution of OPs, for a network that does not contain (does contain) GJ coupling during the first phase of development. The start time of cortical learning is 500 seconds. **B**: Distance from the uniform distribution to the OP distribution for the two networks in A. Smaller values indicate distributions closer to the uniform distribution.

In addition to GJ coupling, the time at which recurrent synapses begin to learn (the start time of the second phase of development) also has an effect on the distribution of OPs. Specifically, the amount of disorder (closeness to a uniform distribution) increases with the start time of recurrent synaptic learning. The intuition is as follows. The sooner that recurrent synapses begin to learn, the less time the feedforward synapses have had to learn, which means that the cortical cells will be less selective by the time that recurrent synapses form. This means that each cell’s OP has not yet been set (the tuning curves are still broad) and communication among cortical cells via synapses will be more likely to influence the final OP formed by each cell. Note that, across all start times of cortical learning, the networks in which GJ-coupling is present during the first phase of development have a more uniform OP distribution than those networks that did not, as shown by the blue curve in Fig 7B. Next, we apply this idea to a cortical network with spatially-restricted synaptic connectivity and analyze the effect of both GJ-coupling during the first phase of development, as well as the effect of timing of the second phase of development on OP organization.

### Realization (iii): recurrent synapse formation (radius of cortical connectivity)

In this section, we introduce spatial restrictions on the potential cortical synaptic connectivity and compare the resulting OP maps across networks that contain GJ-coupling during the first phase of learning and networks that do not. To introduce spatial effects into the model, we draw a fixed radius around each excitatory cortical cell and only allow excitatory synaptic connections from cells within that radius. Note that excitatory to inhibitory, inhibitory to excitatory, and inhibitory to inhibitory synaptic connections still remain all-to-all, with no spatial restrictions. The excitatory to excitatory synaptic strengths are plastic, following the triplet learning rule, while the inhibitory to excitatory synapses are also plastic, following the iSTDP learning rules (recall Fig 2A for a schematic of these learning rules).

Figure 8 shows the development of recurrent synapses onto one sample excitatory neuron in the network. The excitatory recurrent connections within a radius of 4 units are turned on at time t=500 seconds, with initial weights chosen randomly from the interval [0.25, 0.35]*g*_*max*_. As cortical synaptic learning progresses, about half of these excitatory recurrent synapses are potentiated, while half are depressed, as expected and shown in previous sections. The nonzero weights outside of the radius indicate inhibitory synapses onto this excitatory example neuron, which potentiate as the excitatory weights increase to mediate the firing rate of this cell.

**Fig 8.**
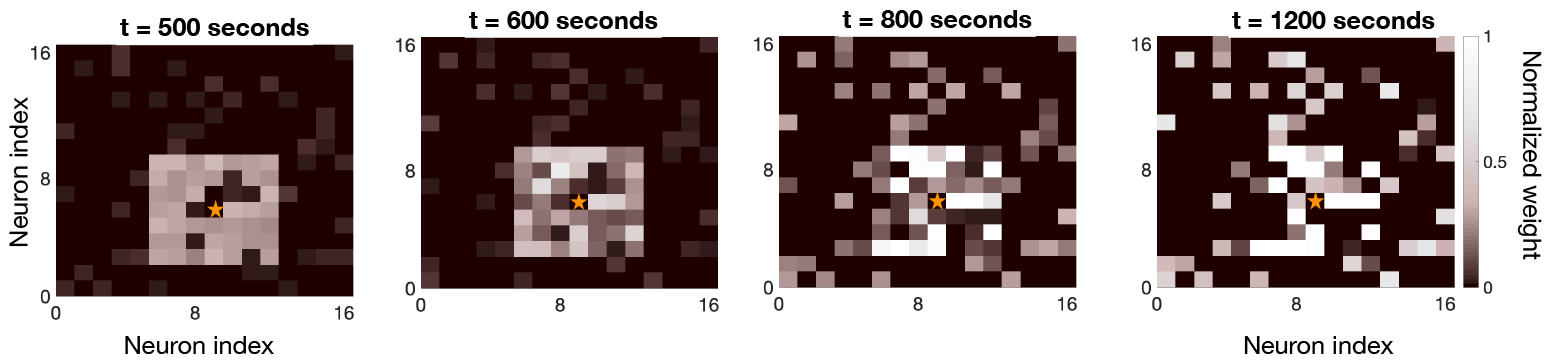
Evolution of the strength of the recurrent synapses onto one sample excitatory cortical cell shown at different time points during the second phase of development. The star indicates the location of the sample neuron. Note that connections outside of the radius of connectivity represent inhibitory synapses onto this excitatory cell.

We anticipate that excitatory cells within the radius of connectivity will follow the results from the all-to-all coupled network in the second network realization. In particular, cells within the radius of connectivity will develop a similar OP in networks that do not contain GJ coupling during the first phase of learning, or in networks for which the second phase of development occurs very soon after the first phase of development (recall Fig 7). On the other hand, if we allow GJ coupling during the first phase of development, or increase the amount of time that feedforward synapses learn without recurrent synapses, we anticipate that the degree of disorder in the OP map will increase. Indeed, Fig 9 demonstrates that our simulations support this hypothesis. The leftmost plot shows the OP map for a network in which the recurrent synapses form at the same time as the LGN synapses. Notice that there are patches of cells with similar OPs, the sizes of which correspond to the radius of connectivity. If we increase the amount of time that the LGN synapses learn without recurrent cortical synapses to 500 seconds (the first phase of development), we observe that the degree of disorder increases until we reach a salt-and-pepper map, see rightmost plots of Fig 9, with the network containing GJ coupling during that first phase of development (top) exhibiting a higher degree of disorder than the network that did not contain GJ coupling during the first phase (bottom).

**Fig 9.**
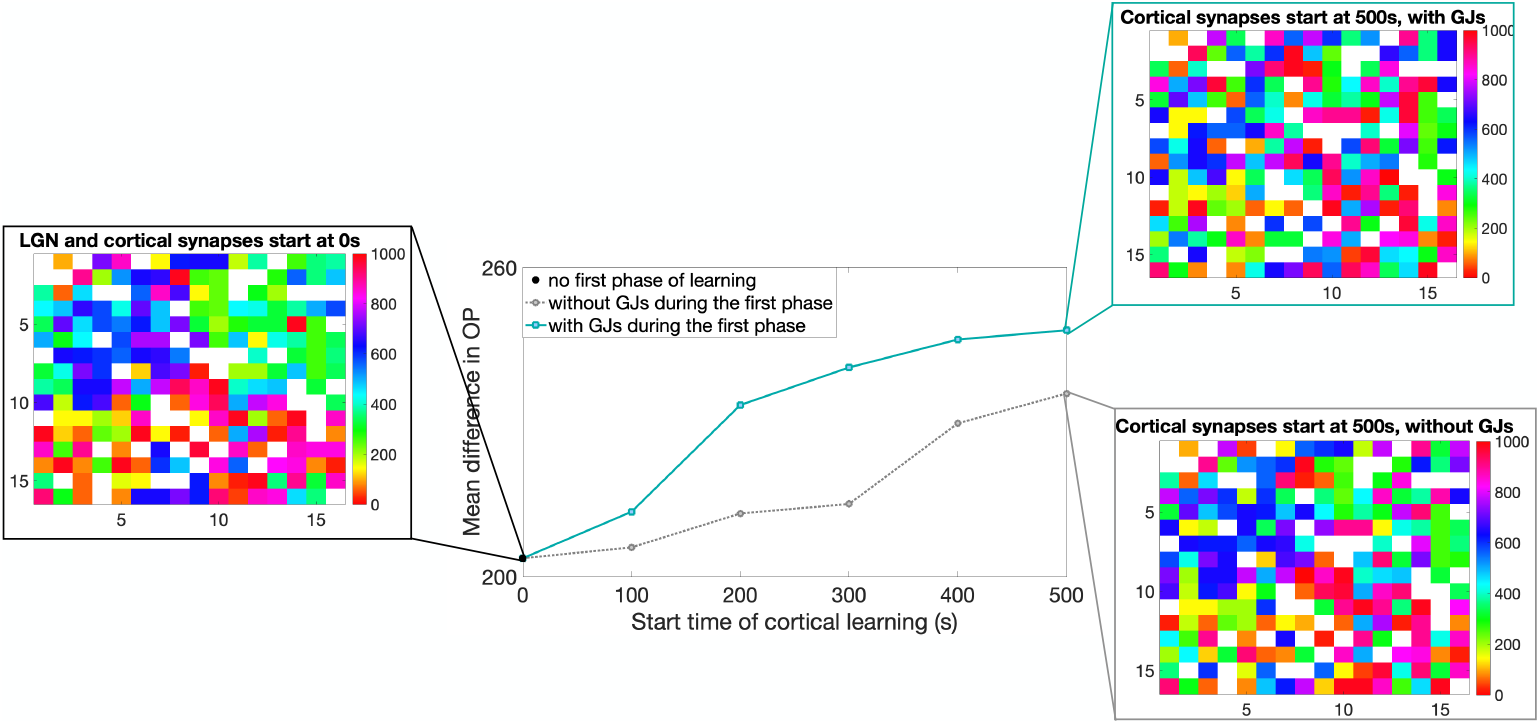
Effect of GJ-coupling and timing of cortical learning on the order of OP maps. The plots shown are OP maps for different types of networks where the color indicates the preference of the cell at that location and the white boxes indicate the inhibitory cells (that do not have a preference). The leftmost map is for a network in which the recurrent synapses form at the same time as the LGN synapses, the top right map is for a network that contains GJ-coupling during the time that LGN synapses are forming (first 500s) and the bottom right map is for a network that does not. The graph in the middle shows the average difference in OP (as defined in the text) for cells within a radius of 4 units, where higher values indicate disorder. The horizontal axis denotes the time at which recurrent synapses within the cortex begin to learn (start time of the second phase of development). After this time, if there were GJs in the network, they are turned off. Note that, for the case of cortical synapses beginning at 0s, there are no GJs in the network by definition since there is no first phase of development.

We quantify the degree of disorder in the OP map by calculating the average difference in OP for each cell within the radius of cortical connectivity. The procedure is as follows. For each excitatory cell, we take the difference between the OP of that cell and the OP of the excitatory cells that are within the radius of connectivity (4 units) and then take the average of those differences. Finally, we take the average of this OP difference over all of the excitatory cells in the network to obtain the measure shown in Fig 9. Notice that larger values of this measure indicate larger differences in OP preference, which corresponds to more disorder. As anticipated, we observe that the degree of disorder in the OP map increases as the start time of the cortical synapses (the length of the first phase of development) increases. The networks in which GJ-coupling is present during the first phase of development follow this same trend as the start time of cortical synapses increases, but also exhibit overall higher levels of disorder than those networks that did not contain GJ coupling; see solid blue curve as compared to dotted gray curve in Fig 9.

We observe that, in these last two realizations of the network model (all-to-all connectivity and radius connectivity), the resulting OP distribution is close to uniform; see Fig 7A and Fig 10. Though each orientation has about equal representation in all example networks, the spatial distribution of the cells with each OP changes drastically across each network (recall the maps plotted in Fig 9).

**Fig 10.**
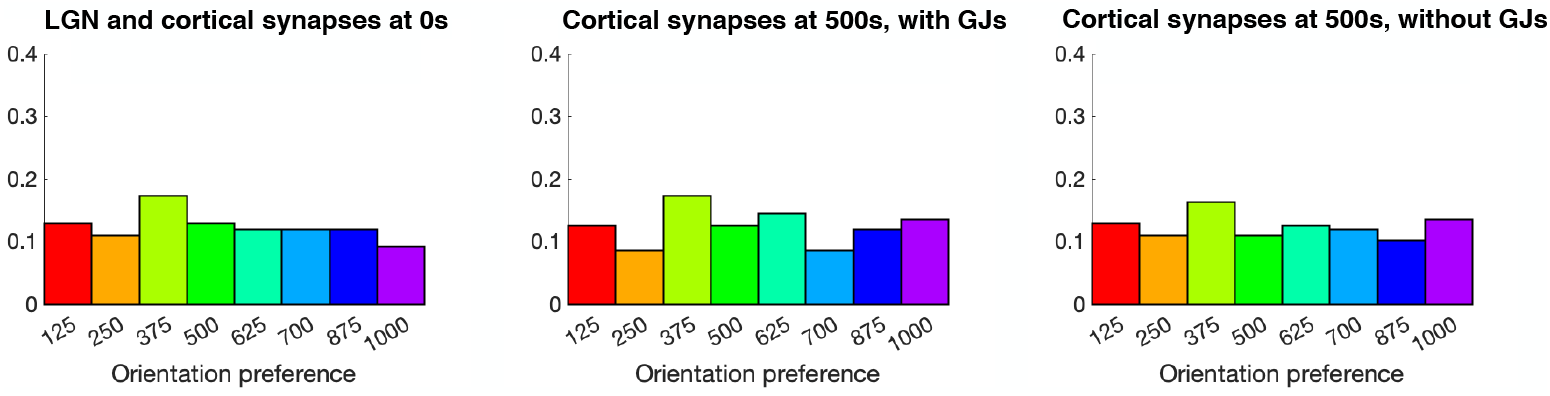
Distribution of OPs for the three example networks shown in Fig 9. The networks, from left to right, are as follows: one in which the recurrent synapses form at the same time as the LGN, one that contains GJs during the 500 seconds that LGN synapses are forming before cortical recurrent connections form, and one that does not contain GJs during the 500 seconds that LGN synapses are forming.

To see if the choice of radius in calculating the difference in OP greatly affects the results shown in Fig 9, we vary the radius used to calculate the difference in OP; see Fig 11. Notice that the measure is low (there is order in the OP map) for small radii, and increases with increasing radius, implying that cells share an OP at small distances, but not at large distances. Importantly, for the networks containing GJ coupling during the first phase of development, and for cases in which the feedforward synapses were allowed to learn for a sufficient amount of time while the GJs are present, there seems to be no order for any value of the radius, see blue and green solid lines in Fig 11. For the same amount of feedforward learning, there is significantly more order for small radii in networks that did not contain GJ coupling, see green dotted curve in Fig 11. This further supports our hypothesis the GJ-coupling during the first phase of development serves to enhance disorder in the OP map.

**Fig 11.**
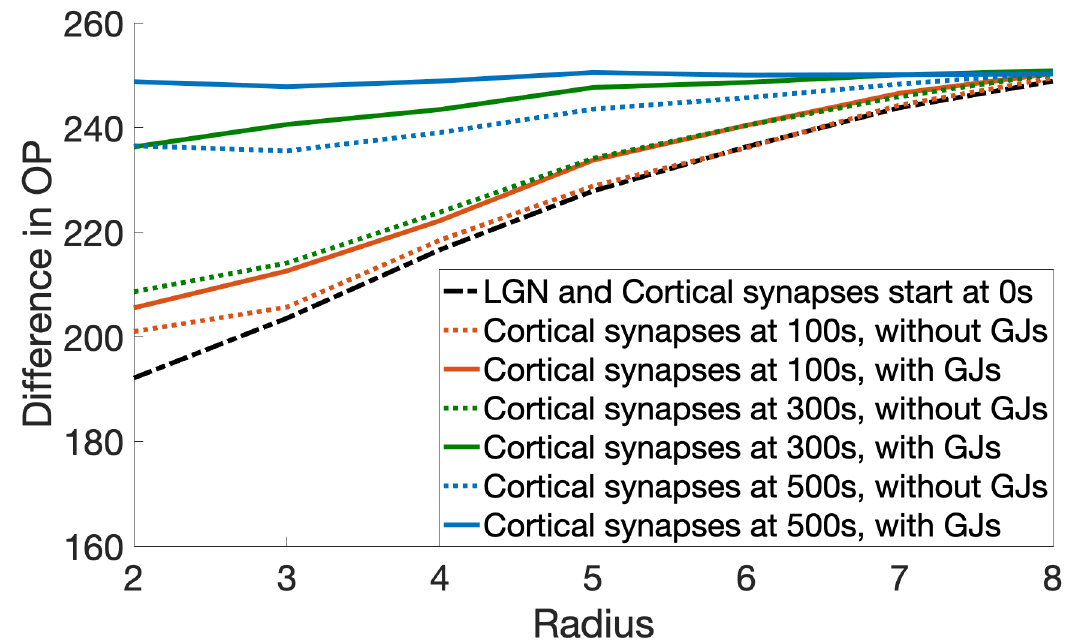
The effect on the average difference in OP with changing radius. The solid lines indicate those networks that contain GJ coupling, while the dotted lines indicate those networks that do not. The colors indicate different start times of the formation of cortical recurrent synapses.

## Discussion

We have created an idealized model to describe the development of synaptic connections both into (LGN) and within (recurrent) the visual cortex of mice during the first two postnatal weeks. The model uses STDP plasticity rules explicitly parametrized for the visual cortex to explore potential mechanisms underlying the formation of ordered or disordered OP maps. We find that GJ coupling during the first postnatal week and relative timing of the recurrent synaptic connections compared with LGN synaptic connections are the two main contributors to order vs. disorder in the adult OP map.

Specifically, we show that the model captures experimentally-measured phenomena such as the preference for cells that were GJ coupled in the first phase of development to share an OP and develop preferentially a bidirectional synapse later in development. In addition to capturing experimentally-observed phenomena, the model also predicts that GJ-coupled cells have a higher firing rate, leading to a faster rate of learning for their LGN synapses when compared to cells that do not have a GJ. We predict that this increased learning rate for sparsely-coupled GJ sister cells, together with the fact that several sets of sister cells are intermingled in the cortex, leads to the formation of a disordered OP map.

We investigate this hypothesis by restricting the recurrent synaptic connections of each cortical cell to a small radius and varying the developmental time at which those recurrent synapses begin learning (which is also the time that GJ-coupling between sister cells disappear). We find that the earlier in developmental time that recurrent synapses are allowed to learn, the more ordered the adult OP map. To understand the influence of GJ-coupling between sister cells, we perform the same analysis for networks in which sister cells were not coupled by GJs in the first phase of development. We find that the timing of recurrent synapses still plays a large role in determining order, but that every network without GJ-coupling in the first phase of development leads to an OP map that exhibits more order than the corresponding network that did have GJ coupling. This leads us to conclude that GJ coupling during the first phase of develop indeed promotes a disordered OP map, but works together with the relative timing of synaptic development from LGN and within the cortex.

The mechanism behind the shared OP of GJ-coupled cells lies in the synchrony (or strongly correlated spike times) induced between the two cells by the GJ. As the LGN synapses form, if the cells are firing synchronously, then those cells will preferentially develop the same set of strengthened LGN synapses, thus forming a similar OP. In our model, sparsity of GJ coupling between the sister cells is essential for this synchrony to occur, and consequently for the shared OP of GJ-coupled cells. When cells are coupled with a probability of 5%, each cell is coupled to an average of 1.5 other cells, leading to isolated pairs or triplets of GJ-coupled cells. As the coupling percentage increases, there are no longer isolated pairs or triplets of cells; rather, each cell may be coupled to several different groups of GJ-coupled sister cells, leading to an overall desynchronization of those cells; see S1 Fig. We note, however, that though the shared OP of GJ-coupled cells is diminished with an increase in GJ-coupled probability, all other properties of OP-map development, such as the discussion of order vs. disorder, do not rely on this assumption; see S2 Fig. In addition, though experimental measurements of the GJ-coupling probability between sister cells during the first postnatal week is about 28%, this was specifically measured for isolated pairs of radially-aligned sister cells [7]. In this work, we are interested in sister cells that are GJ-coupled laterally (within the layer), which hasn’t explicitly been measured.

Finally, we note that the model developed in this work is highly idealized, especially in its size, spatial structure, and LGN input organization. Regardless, the model reproduces experimentally-measured properties of GJ-coupled cells and uses these properties to propose two mechanisms affecting the formation of salt-and-pepper OP maps in the mouse V1: the presence of GJs during the first postnatal week and the relative timing of cortical synapse formation to the timing of LGN synapse formation. In the future, we plan to develop a comprehensive large-scale model of the developing visual cortex, including realistic LGN input and spatial organization of the cortex, to further test the hypotheses presented in this work.

## Supporting information

**S1 Fig.**
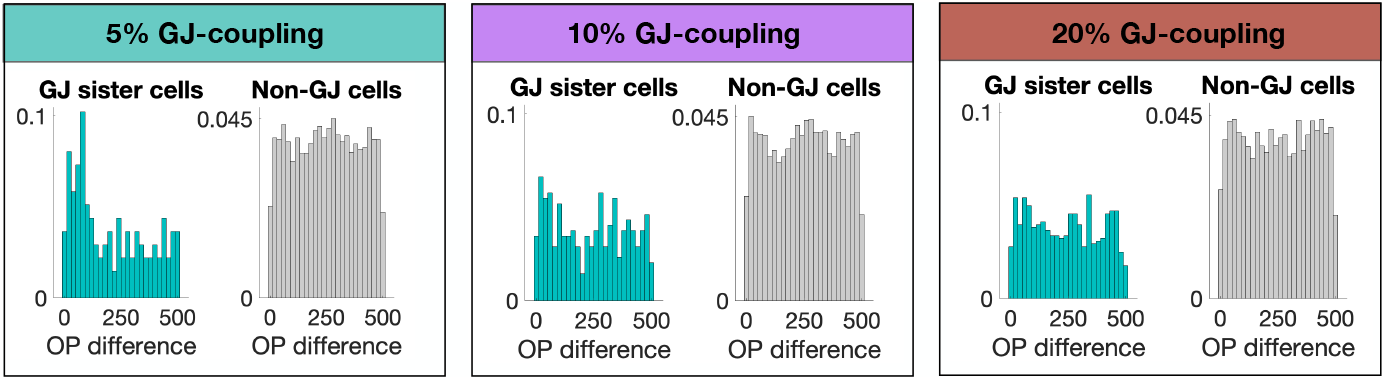
Increasing the number of GJ connections per neuron disrupts the OP-sharing preference of GJ-coupled cells due to less synchronous firing. In the case of a low coupling percentage, the average number of GJ connections per sister cell is 1.5, leading to more cases of isolated GJ-coupled pairs or triplets. For higher percentages, the average number of GJ connections per sister cell increases to 3.5 for 10% GJ coupling and 6.8 for 20% GJ coupling (note that there are only about 30 cells in each sister group on average). This leads to larger networks of GJ-coupled cells, more instances of cells that are GJ-coupled to more than one GJ-coupled group, and less synchrony of the GJ-coupled cells. A direct effect of this decrease in synchrony is a disruption in the OP-sharing property of GJ-coupled cells.

**S2 Fig.**
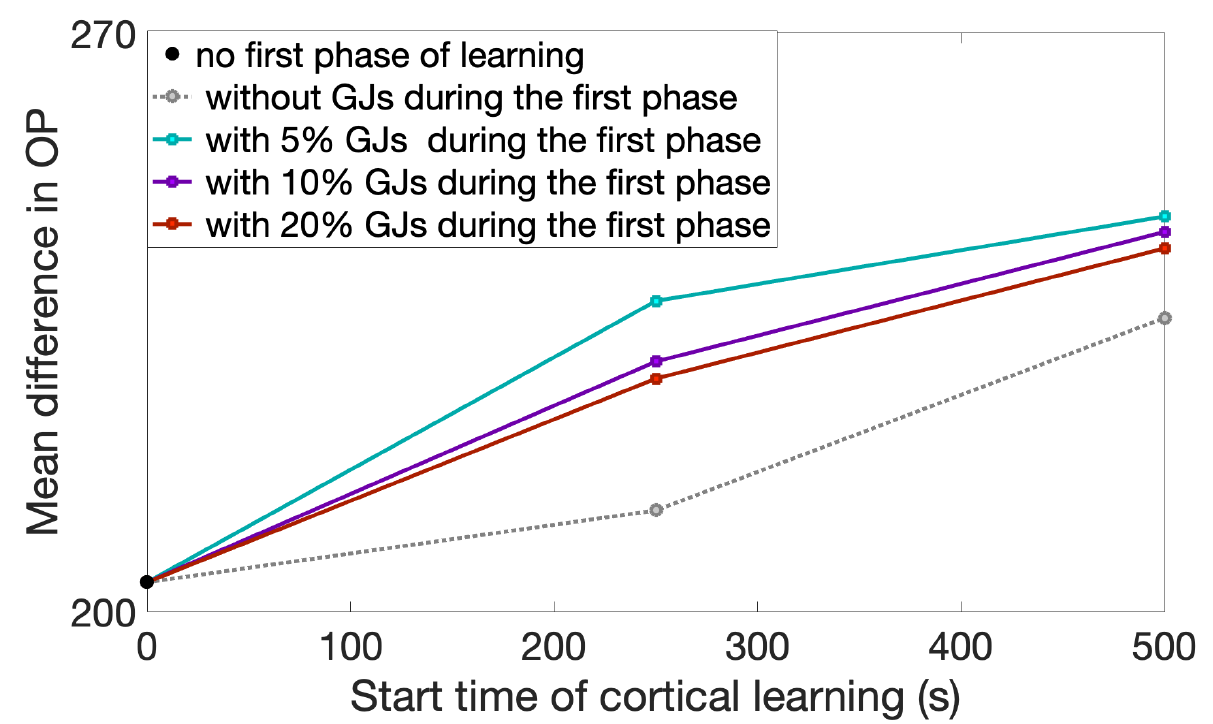
The OP-map formation is largely unchanged by increasing the GJ-coupling probability. No GJ-coupling during the first phase of development leads to an ordered map, while the inclusion of GJs leads to disorder, independent of the GJ connectivity percentage. Note that we only show results for greater than 250 seconds of cortical learning to ensure the GJ-coupling has time to affect the dynamics.

## Acknowledgments

We would like to thank Dr. Barry Connors for meaningful and helpful conversation early in the work.

## References

1. M. V. L. Bennett and R. S. Zukin. Electrical coupling and neuronal synchronization in the mammalian brain. Neuron, 41(4):495–511, Feb 2004.

2. G. Tamas, E. H. Buhl, A. Lorincz, and P. Somogyi. Proximally targeted gabaergic synapses and gap junctions synchronize cortical interneurons. Nat. Neurosci., 3(4):366–371, Apr 2000.

3. Y. Wang, A. Barakat, and H. Zhou. Electrotonic coupling between pyramidal neurons in the neocortex. PLoS One, 5(4):e10253, 2010.

4. Lopez-Bendito G, Molnar Z. Thalamocortical development: how are we going to get there? Nat Rev Neurosci. 2003;4(4):276–289. doi:10.1038/nrn1075.

5. Yu YC, Bultje RS, Wang X, Shi SH. Specific synapses develop preferentially among sister excitatory neurons in the neocortex. Nature. 2009;458(7237):501–504. doi:10.1038/nature07722.

6. Li M, Cui Z, Niu Y, Liu B, Fan W, Yu D, et al. Synaptogenesis in the developing mouse visual cortex. Brain Res Bull. 2010;81(1):107–113. doi:10.1016/j.brainresbull.2009.08.028.

7. Yu YC, He S, Chen S, Fu Y, Brown KN, Yao XH, et al. Preferential electrical coupling regulates neocortical lineage-dependent microcircuit assembly. Nature. 2012;486(7401):113–117. doi:10.1038/nature10958.

8. Pangratz-Fuehrer S, Hestrin S. Synaptogenesis of Electrical and GABAergic Synapses of Fast-Spiking Inhibitory Neurons in the Neocortex. The Journal of neuroscience : the official journal of the Society for Neuroscience. 2011;31:10767–75. doi:10.1523/JNEUROSCI.6655-10.2011.

9. Espinosa JS, Stryker MP. Development and plasticity of the primary visual cortex. Neuron. 2012;75(2):230–249. doi:10.1016/j.neuron.2012.06.009.

10. Siegel F, Heimel JA, Peters J, Lohmann C. Peripheral and central inputs shape network dynamics in the developing visual cortex in vivo. Curr Biol. 2012;22(3):253–258. doi:10.1016/j.cub.2011.12.026.

11. Ko H, Cossell L, Baragli C, Antolik J, Clopath C, Hofer SB, et al. The emergence of functional microcircuits in visual cortex. Nature. 2013;496(7443):96–100. doi:10.1038/nature12015.

12. Song S, Abbott LF. Cortical development and remapping through spike timing-dependent plasticity. Neuron. 2001;32(2):339–350.

13. Li Y, Lu H, Cheng PL, Ge S, Xu Ht, Shi SH, et al. Clonally Related Visual Cortical Neurons Show Similar Stimulus Feature Selectivity. Nature. 2012;486:118–21. doi:10.1038/nature11110.

14. Nadarajah B, Jones AM, Evans WH, Parnavelas JG. Differential expression of connexins during neocortical development and neuronal circuit formation. J Neurosci. 1997;17(9):3096–3111.

15. Ko H, Hofer S, Pichler B, A Buchanan K, Sjöström P, Mrsic-Flogel T. Functional specificity of local connections in neocortical networks. Nature. 2011;473:87–91. doi:10.1038/nature09880.

16. K. M. Hagihara, T. Murakami, T. Yoshida, Y. Tagawa, and K. Ohki. Neuronal activity is not required for the initial formation and maturation of visual selectivity. Nat Neurosci, 18(12):1780–1788, Dec 2015.

17. Ohtsuki G, Nishiyama M, Yoshida T, Murakami T, Histed M, Lois C, et al. Similarity of visual selectivity among clonally related neurons in visual cortex. Neuron. 2012;75(1):65–72. doi:10.1016/j.neuron.2012.05.023.

18. Magavi S, Friedmann D, Banks G, Stolfi A, Lois C. Coincident generation of pyramidal neurons and protoplasmic astrocytes in neocortical columns. J Neurosci. 2012;32(14):4762–4772. doi:10.1523/JNEUROSCI.3560-11.2012.

19. Tan SS, Breen S. Radial mosaicism and tangential cell dispersion both contribute to mouse neocortical development. Nature. 1993;362(6421):638–640. doi:10.1038/362638a0.

20. Torii M, Hashimoto-Torii K, Levitt P, Rakic P. Integration of neuronal clones in the radial cortical columns by EphA and ephrin-A signalling. Nature. 2009;461(7263):524–528. doi:10.1038/nature08362.

21. Lewis TJ, Rinzel J. Dynamics of spiking neurons connected by both inhibitory and electrical coupling. J Comput Neurosci. 2003;14(3):283–309.

22. Pfister JP, Gerstner W. Triplets of spikes in a model of spike timing-dependent plasticity. J Neurosci. 2006;26(38):9673–9682. doi:10.1523/JNEUROSCI.1425-06.2006.

23. Bi GQ, Poo MM. Synaptic modifications in cultured hippocampal neurons: dependence on spike timing, synaptic strength, and postsynaptic cell type. J Neurosci. 1998;18(24):10464–10472.

24. Sjostrom PJ, Turrigiano GG, Nelson SB. Rate, timing, and cooperativity jointly determine cortical synaptic plasticity. Neuron. 2001;32(6):1149–1164. doi:10.1016/s0896-6273(01)00542-6.

25. Zenke F, Hennequin G, Gerstner W. Synaptic plasticity in neural networks needs homeostasis with a fast rate detector. PLoS Comput Biol. 2013;9(11):e1003330. doi:10.1371/journal.pcbi.1003330.

26. S. Song, K. D. Miller, and L. F. Abbott. Competitive hebbian learning through spike-timing-dependent synaptic plasticity. Nat Neurosci, 3(9):919–926, Sep 2000.

27. Shen J, Colonnese MT. Development of Activity in the Mouse Visual Cortex. J Neurosci. 2016;36(48):12259–12275. doi:10.1523/JNEUROSCI.1903-16.2016.

28. Vogels TP, Sprekeler H, Zenke F, Clopath C, Gerstner W. Inhibitory plasticity balances excitation and inhibition in sensory pathways and memory networks. Science. 2011;334(6062):1569–1573. doi:10.1126/science.1211095.

